# Human iPSC-derived astrocytes transplanted into the mouse brain display three morphological responses to amyloid-β plaques

**DOI:** 10.1101/2020.11.19.389023

**Authors:** Pranav Preman, Julia TCW, Sara Calafate, An Snellinx, Maria Alfonso-Triguero, Nikky Corthout, Sebastian Munck, Dietmar Rudolf Thal, Alison M Goate, Bart De Strooper, Amaia M Arranz

## Abstract

**Background:** Increasing evidence for a direct contribution of astrocytes to neuroinflammatory and neurodegenerative processes causing Alzheimer’s disease comes from molecular studies in rodent models. However, these models may not fully recapitulate human disease as human and rodent astrocytes differ considerably in morphology, functionality, and gene expression.

**Methods:** To address these challenges, we established an approach to study human astroglia within the context of the mouse brain by transplanting human induced pluripotent stem cell (hiPSC)-derived glia progenitors into neonatal brains of immunodeficient mice.

**Results:** Xenografted (hiPSC)-derived glia progenitors differentiate into astrocytes that integrate functionally within the mouse host brain and mature in a cell-autonomous way retaining human-specific morphologies, unique features and physiological properties. In Alzheimer’s chimeric brains, transplanted hiPSC-derived astrocytes respond to the presence of amyloid plaques with various morphological changes that seem independent of the *APOE* allelic background.

**Conclusion:** In sum, this chimeric model has great potential to analyze the role of patient-derived and genetically modified astroglia in Alzheimer’s disease.

## BACKGROUND

Astrocytes are essential to maintain the homeostasis of the brain, provide trophic support, stimulate synaptogenesis and neurotransmission, and regulate blood-brain-barrier permeability (1,2). Impaired astroglial function contributes to neurological and neurodegenerative disorders including Alzheimer’s disease (AD) (3–8). Genome-wide association studies (9,10) show that genetic risk of AD is also associated with genes mainly expressed in astroglia such as Clusterin (*CLU*), Fermitin family member 2 (*FERMT2*) and Apolipoprotein E (*APOE*) (11), highlighting the potential importance of these cells in the disease. Different types of astroglial pathology have been described in the AD brain (12–14). Among those, hypertrophic (15), quiescent and degenerating morphologies (16,17) were found.

Transgenic models have provided invaluable tools to study the role of astroglia in AD (18–21). However, these models of AD might insufficiently mimic the human disease, as there are major differences between rodent and human astrocytes. Morphologically, human astrocytes are larger and more complex, having around 10 times more processes than their rodent counterparts (22). Molecularly, human astrocytes and mouse astrocytes display different, although overlapping, gene expression profiles (11). Functionally, human astrocytes propagate calcium waves four-fold faster than rodent ones (11,22,23), and human and mouse astrocytes show very different responses when exposed to inflammatory stimuli (24,25).

The ability to generate induced pluripotent stem cells (iPSCs) from patients and differentiate them into astrocytes and other CNS cell types has generated exciting opportunities to examine AD associated phenotypes *in vitro* (39) and unravel the contribution of astroglial risk genes to AD (26–29). Yet, human iPSC (hiPSC)-derived astrocytes grown in culture lack essential components present in the brain which can induce altered phenotypes and gene expression signatures significantly different from that of primary resting astroglia in the brain (11,30). Therefore, it has proved challenging to advance understanding of human astroglial function in AD.

To address these challenges, we aimed at developing a chimeric model that allows studying hiPSC-derived astrocytes in an *in vivo* AD context. We and others have generated chimeric models to study AD by transplanting human PSC-derived neurons or microglia into the brains of immunodeficient AD mice and wild-type littermates (31–33). These models revealed that human neurons and microglia transplanted into the mouse brain respond to pathology differently than their murine counterparts, showing specific vulnerability and transcriptional signatures when exposed to amyloid-β (Aβ) (31,32). Moreover, human glia chimeric mice have been generated by Goldman and collaborators to investigate the function of engrafted human glia, mainly NG2 cells and lower proportions of oligodendrocytes and astrocytes, in disease relevant conditions such as Huntington disease, Schizophrenia or hypomyelination (34–36). Yet, to date no studies have analyzed the phenotype and functional responses of xenografted human astrocytes exposed to Aβ and AD-associated pathology *in vivo*.

We established here a chimeric model to investigate survival, integration, properties and responses to Aβ species of human astrocytes expressing *APOE* ɛ3 (E3) vs *APOE* ɛ4 (E4) variants. We document here engraftment of astrocytes that integrate in a functional way in the mouse host brain and display human-specific morphologies and properties. When transplanted human astrocytes are exposed to Aβ plaques, they display hypertrophic and atrophic responses similar to the ones seen in AD patients’ brains (12,16,17). Our results validate the use of chimeric mice as a potential powerful tool for studying astrocyte contribution to AD. We also discuss one of the major hurdles to fully capture the strength of this approach, which is, in our hands, the variable and often low degree of chimerism obtained with human astrocytes from different hiPSC lines after several months of transplantation.

## METHODS

### Generation of isogenic CRISPR/Cas9 gene-edited hiPSCs

Eight hiPSC lines were generated from three *APOE* ɛ4 carriers diagnosed with AD (Table 1) as described previously by the ‘CORRECT’ scarless gene-editing method (37). The correct *APOE* sgRNA sequence orientation was confirmed by Sanger sequencing and CRISPR/Cas9-*APOE* sgRNA plasmid cleavage efficiency was determined using the Surveyor mutation detection kit in 293T cells. The single-strand oligo-deoxynucleotide (ssODN) was designed to convert *APOE* ɛ4 to *APOE* ɛ3 with a protospacer adjacent motif (PAM) silent mutation to prevent recurrent Cas9 editing. hiPSCs (70-80% confluent) dissociated by Accutase supplemented with 10 μM Thiazovivin (Tzv) (Millipore), were harvested (200 × g, 3 min), and electroporated (Neon^®^, ThermoFisher) according to the manufacturer’s instructions. In brief, cells resuspended in 10μl Neon Resuspension Buffer R, 1μg CRISPR/Cas9-*APOE* sgRNA plasmid and 1μl of 10μM of ssODN were electroporated plated on Matrigel-coated plates in mTeSR media with 10 μM Tzv for 72h. GFP-expressing hiPSC were isolated by FACS (BD FACSAria). Sorted single cells were suspended in mTeSR with Tzv and plated into 96 well plates containing MEFs (4,000 cells/well). Clones were expanded and transferred to a replicate plate for gDNA isolation and Sanger sequencing to identify genome edited clones.

**Table 1.**
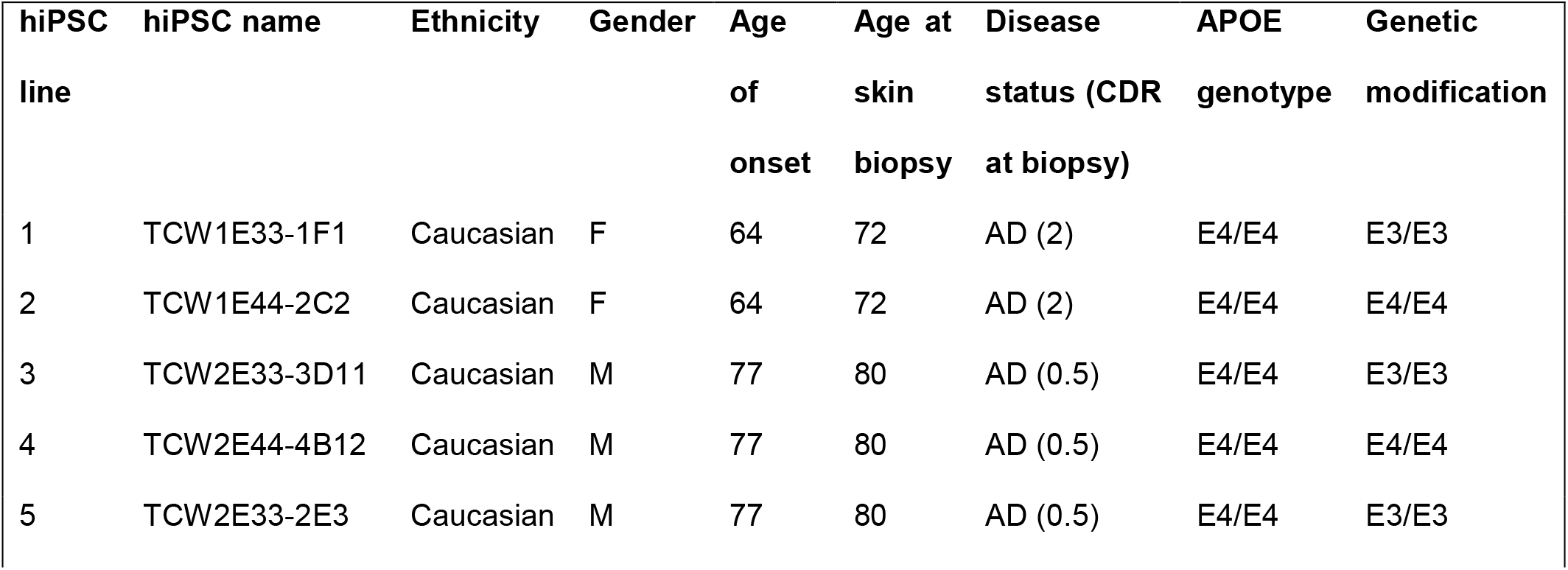
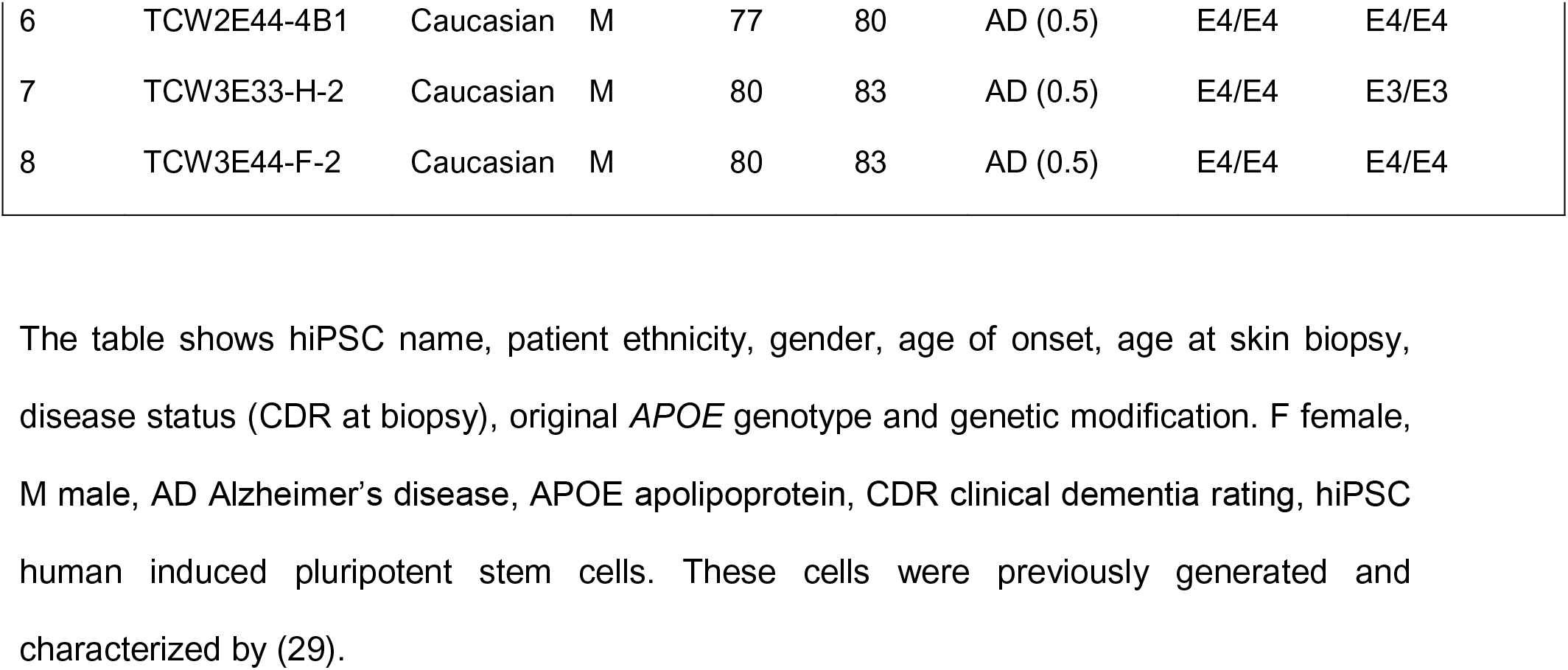
Information on the hiPSC lines.

### Karyotyping

Karyotyping was performed by Wicell Cytogenetics (Madison, WI). Karyotypes are shown in Additional file 2, Figure S1.

### Generation of reporter hiPSC-astrocytes

The consent for reprogramming human somatic cells to hiPSC was carried out on ESCRO protocol 19-04 at Mount Sinai (J.TCW.). hiPSCs maintained on Matrigel (Corning) in mTeSR1 (StemCell Technologies) supplemented with 10 ng/ml FGF2 StemBeads (StemCultures) were differentiated to neural progenitor cells (NPCs) by dual SMAD inhibition (0.1μM LDN193189 and 10μM SB431542) in embryoid bodies (EB) media (DMEM/F12 (Invitrogen, 10565), 1x N2 (Invitrogen, 17502-048), and 1x B27-RA (Invitrogen, 12587-010)). Rosettes were selected at 14 DIV by Rosette Selection Reagent (StemCell Technologies) and patterned to forebrain NPCs with EB media containing 20ng/ml FGF2 (Invitrogen). NPCs (CD271^−^/CD133^+^) were enriched by magnetic activated cell sorting (Miltenyi Biotec) (38) and validated immunocytochemically using SOX2, PAX6, FoxP2 and Nestin (Additional file 1, Table S1). Dissociated single cell forebrain NPCs were plated 1,000,000 cells/well on 12 well plates and transfected with lentiGuide-tdTomato (Addgene #99376) plasmid and selected by hygromycine. Pure fluorescent expressing NPCs were plated at low density (15,000 cells/cm^2^) on matrigel coated plates and differentiated to astrocytes in astrocyte medium (ScienCell, 1801) as described (39). Cells were cultured and harvested as astroglia progenitors at DIV 40-44, validated immunocytochemically and/or by FACS for the astrocyte-specific markers and used for subsequent experiments.

### AD and WT Immunodeficient Mice

Mice were generated as described previously (31). Briefly, APP PS1 tg/wt mice (expressing KM670/671NL mutated APP and L166P mutated PS1 under the control of the Thy1.2 promoter1.1) (40) were crossed with the immunodeficient NOD-SCID mice (NOD.CB17-Prkdc^scid^) that carry a single point mutation in the Prkdc gene (41). APP PS1 tg/wt Prkdc^scid/+^ mice from the F1 generation were crossed with NOD-SCID mice to generate APP PS1 tg/wt Prkdc^scid/scid^ immunodeficient mice. APP PS1 tg/wt Prkdc^scid/scid^ mice were subsequently crossed with NOD-SCID mice to generate either APP PS1 tg/wt Prkdc^scid/scid^ (AD mice) or APP PS1 wt/wt Prkdc^scid/scid^ (WT mice) used for transplantations. Mice were housed in IVC cages in a SPF facility; light/dark cycle and temperature were always monitored. After weaning, no more than five animals of the same gender were kept per cage. Genotyping was done as previously described (31). Transplantation experiments were performed in both male and female littermates at P0-P4. Mouse work was performed in accordance with institutional and national guidelines and regulations, and following approval of the Ethical Committee of the KUL. All experiments conform to the relevant regulatory standards.

### Intracerebral Grafting

Grafting experiments of hiPSC-derived glial progenitors using neonatal APP PS1 tg/wt NOD-SCID (AD mice) and APP PS1 wt/wt NOD-SCID (WT mice) at postnatal days P0-P4 were performed as described previously (31) with some modifications. Briefly, hiPSC-derived glia progenitor cells at DIV 44 were enzymatically dissociated, supplemented with HB-EGF (100-47, Peprotech) and RevitaCell (A2644501, ThermoFisher) and injected into the frontal cortex of AD or WT mice. The pups were anesthetized by hypothermia and about 200,000 cells were injected with Hamilton syringes into the forebrain at two locations: 1 mm posterior Bregma, 1.5 mm bilaterally from the midline and 1.2 mm from the pial surface. Transplanted pups were returned to their home cages until weaning age.

### Electrophysiological Characterization of Human Glia in Chimeric Mice

Four to five month-old WT mice were anesthetized with isoflurane and decapitated. Acute 300 μm-thick coronal slices were cut on a Leica VT1200 vibratome in a sucrose-based cutting solution consisting of (mM): 87 NaCl, 2.5 KCl, 1.25 NaH2PO4, 10 glucose, 25 NaHCO3, 0.5 CaCl2, 7 MgCl2, 75 sucrose, 1 kynurenic acid, 5 ascorbic acid, 3 pyruvic acid (pH 7.4 with 5% CO2/ 95% O2). Slices were allowed to recover at 34°C for 45 minutes and maintained at room temperature (RT) in the same solution for at least 30 minutes before using. During recordings, slices were submerged in a chamber (Warner Instruments) perfused with 3-4mL/min artificial cerebrospinal fluid (ACSF) consisting of (mM): 119 NaCl, 2.5 KCl, 1 NaH2PO4, 26 NaHCO3, 4 MgCl2, 4 M CaCl2, 11 glucose at pH 7.4 with 5% CO2/ 95% O2. Recordings were done at 34°C. hiPSC-astrocytes were identified based on the td-Tomato fluorescence with a 40x objective in an epifluorescent microscope (Zeiss Axio Examiner.A1). Whole-cell current clamp recordings were made from 17 hiPSC-astrocytes (hiPSC lines #1 to #4, n=6 mice) with borosilicate glass recording pipettes (resistance 3-6MΩ). Pipettes were pulled on a horizontal micropipette puller (Sutter P-1000) and filled with a K-gluconate based internal medium consisting of (mM): 135 K-Gluconate, 4 KCl, 2 NaCl, 10 HEPES, 4 EGTA, 4 MgATP, 0.3 NaATP (pH 7.25). To post-hoc identify the patched astrocyte and analyze its potential to form gap-junctions, 40 μM Alexa Fluor hydrazide dye 488 (Invitrogen) was included in the internal medium. Current steps of incrementing 20 pA were injected starting from 50 pA up to 150 pA. Resting membrane potential was calculated using Clampfit 10.7 (Axon Instruments). Currents were sampled at 20 kHz and stored after 3 kHz low-pass Bessel filtering. The data was low-pass filtered at 1 kHz (Molecular devices DigiData 1440A and Multiclamp 700B). Pipette series resistance and membrane holding current were monitored throughout all recordings to ensure stability of the recording.

### Immunofluorescence (IF) in Chimeric Mice

For IF analysis, mice were anesthetized with CO2 and perfused with phosphate-buffered saline followed by 4% paraformaldehyde solution. The brain was then removed, post-fixed in the same fixative overnight to 48 hr and cut into 40 μm slices on a Leica VT1000S vibratome. IF on grafted brains was performed as described previously (31) using primary and secondary antibodies (Additional file 1, Table S1). Antigen retrieval was performed by microwave boiling the slides in 10mM tri-Sodium Citrate buffer pH 6.0 (VWR). Aβ plaques were detected by staining with Thioflavin (SIGMA). Briefly, for Thioflavin staining brain sections were incubated with a filtered 0.05% aqueous Thioflavin-S (SIGMA) solution in 50% ethanol for 5 min at RT and rinsed gradually with 70%, 95% ethanol and water. Nuclei staining was performed using a specific anti-human Nuclear Antigen antibody (hNuclei) (Additional file1, Table S1), the pan-nuclear staining TOPRO3 (Invitrogen), or DAPI (SIGMA). The sections were mounted with Glycergel (DAKO). Confocal images were obtained using a Nikon Ti-E inverted microscope equipped with an A1R confocal unit driven by NIS (4.30) software. The confocal was outfitted with 20x (0.75 NA), 40x oil (1.4 NA) and 60x oil (1.4 NA) objectives lenses. For excitation 405 nm, 488 nm, 561 nm, 638 nm laser lines were used.

### Quantification and Statistical Analysis

Morphometry and measurements were performed with Fiji/ImageJ software on animals at five months after transplantation. At least 4-5 different coronal brain sections comprising the transplanted astrocytes and the mouse host tissue were included per animal. Immunofluorescence (IF) sections were imaged by confocal microscopy (Nikon Ti-E inverted microscope) using a 20x (0.75 NA) objective lens to image Z-stacks (8-10 optical sections with a spacing of 1 μm). All images were acquired using identical acquisition parameters as 16-bit, 1024×1024 arrays. Maximum intensity projections and threshold were applied using Fiji/ImageJ to isolate specific fluorescence signals.

For analyses of **cell integration**, brains were sectioned and stained with the antibodies against RFP and hNuclei (human Nuclear antigen). The number of hNuclei+ and RFP+ cells was counted manually on IF images of astrocytes derived from the eight hiPSC lines used on the study (#1 to #8, Table 1). Final counts were corrected for series number (1:6) to get an estimate of the total number of hNuclei+ and RFP+ cells per animal (Additional file 2, Figure S1d).

For analyses of **cell identity**, brains were sectioned and stained with the following antibodies: RFP and hNuclei (human Nuclear antigen), GFAP (astroglia marker), NeuN (neuronal marker) or APC (marker of oligodendrocytes). Results are shown for four hiPSC lines (#1, #2, #7 and #8, Table 1). Total percentages of RFP+ cells co-localizing with GFAP (n=14 mice), hNuclei (n=15 mice), NeuN or APC (n=9 mice each) were manually determined on IF images using Fiji/ImageJ. Data are represented as mean ± SEM. Statistical analyses were done with Student’s t test (Fig. 1 and Additional file 3, Figure S2).

**Fig. 1.**
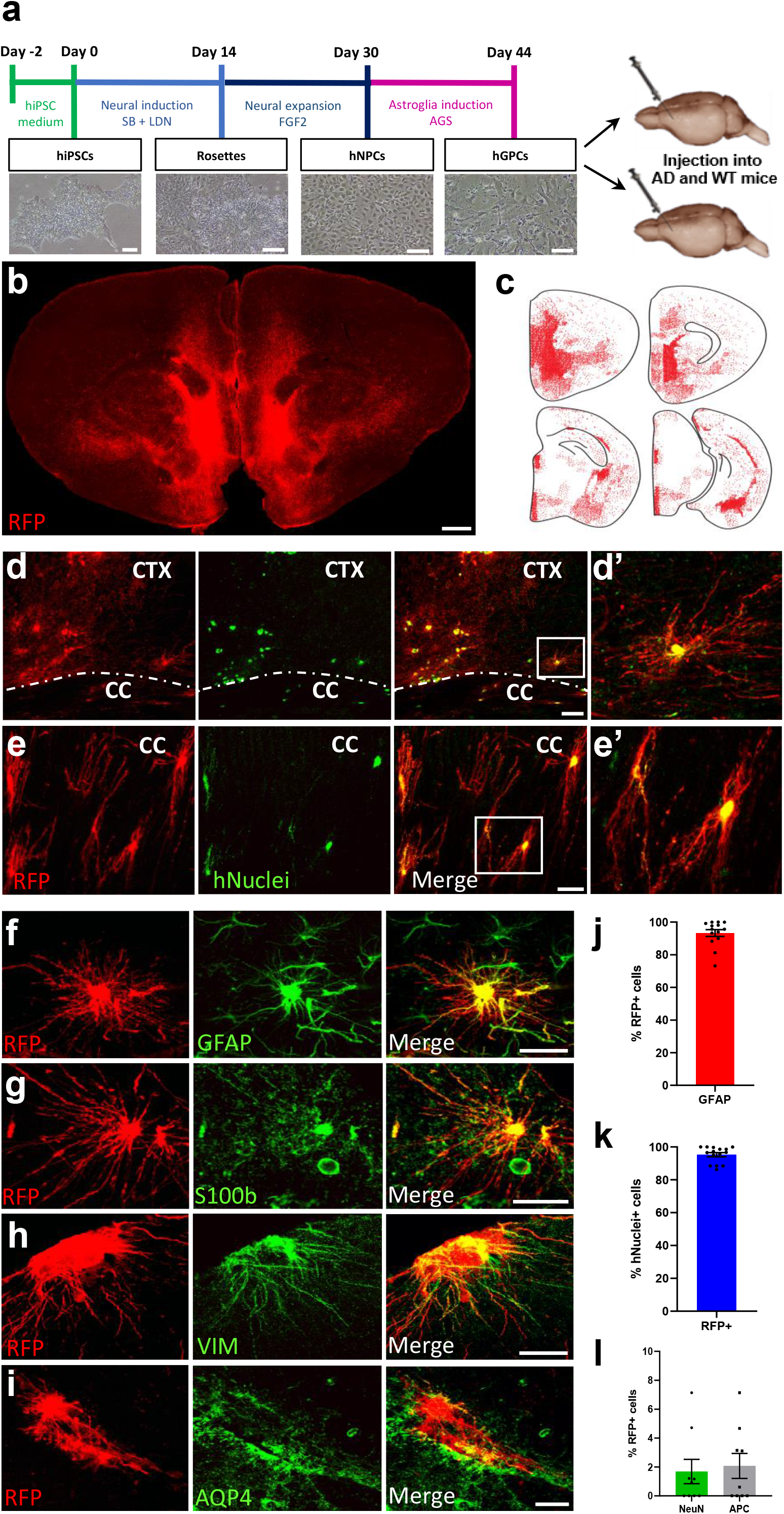
hiPSC-glia progenitors engraft the mouse brain and differentiate into astrocytes. **(a)** Schematics of the differentiation and transplantation procedures. hiPSCs: human induced pluripotent stem cells, NPCs: neural progenitor cells, GPCs: glia progenitor cells, SB: SB431542, LDN: LDN193189, FGF2: fibroblast growth factor 2, AGS: astrocyte growth supplement. Scale bars: 100 μm. **(b)** RFP staining (red) shows the distribution of hiPSC-derived astrocytes on a coronal brain section of a chimeric mouse at five months after transplantation. Scale bar: 200 μm. **(c)** Dot map displaying the widespread distribution of the hiPSC-derived astrocytes (RFP, red) in four coronal sections of this mouse brain. **(d-e)** RFP (red) and hNuclei (green) expressing hiPSC-astrocytes depict a complex fine structure in the cortex (CTX) and corpus callosum (CC) of chimeric mice. Scale bars: 50 μm (d), 25 μm (e). **(d’-e’)** Enlarged images of the inserts in d and e. **(f-i)** Engrafted hiPSC-astrocytes (RFP+, red) express GFAP (f), S100b (g), Vimentin (h) and AQP4 (i) (green) five months after transplantation. Scale bars: 25 μm. **(j)** Percentage of RFP+ cells expressing GFAP (n=14 mice). **(k)** Percentage of hNuclei+ cells expressing RFP (n=15 mice). **(l)** Percentage of RFP+ cells expressing NeuN and APC (n=9 mice). Data are represented as mean ± SEM

To analyze the **morphological subtypes of hiPSC-astrocytes**, brains were sectioned and stained with antibodies against RFP and hNuclei (human Nuclear antigen) and morphometry analyses were manually performed on IF images using Fiji/ImageJ. Results are shown for two hiPSC lines (#1 and #2, Table1) in WT mice (n=9). Data are represented as mean ± SEM (Fig. 3).

For quantification of the average **cell area**, brains were stained with RFP and GFAP, and the NIS-elements software was used (version 5.21.01 build 1483, Nikon Instruments). All the z-stacks were first denoised (denoise.ai tool) and then projected on a 2D image using an extended focus operation (EDF, zero-based, balanced). The resulting 2D image was used for further quantification with a General Analysis (GA3) protocol. In short, to count the number of cells, a spot detection approach was used (average size 11 μm). For detection of the cell area, we first applied a rolling ball filter (6 μm) and, consequently, a thresholding step. Both the settings for the threshold and the spot detection were adjusted per image to compensate for differences in intensity due to a change of acquisition parameters. Results are shown for four hiPSC lines (#1, #2, #3 and #4, Table1) in WT mice (n=12). Data are represented as mean ± SEM. Statistical analysis was done with Student’s t test (Fig. 3).

To analyze the **morphological responses to Aβ plaques**, brains were sectioned and stained with RFP and Thioflavin and morphometry analyses were manually performed on IF images using Fiji/ImageJ. Results are shown for two hiPSC lines (#1 and #2, Table1) in AD mice (n=7). Data are represented as mean ± SEM. Statistical analysis was performed with Chi-square t test (Fig. 5).

### Neuropathology on Human Brain Samples

Brain tissue samples from 4 AD, 5 pre-AD and 3 non-demented control patients were included in this study (Table 2). The autopsies were performed with informed consent in accordance with the applicable laws in Belgium (UZ Leuven) and Germany (Ulm, Bonn and Offenbach). The use of human tissue samples for this study was approved by the UZ Leuven ethical committee (Leuven, Belgium). Brain tissues were collected as described in previous studies (42) with an average post-mortem interval (PMI) of 48 h. Briefly, after autopsy, the brains were fixed in 4% aqueous solution of formaldehyde for 2–4 weeks. Samples of the anterior entorhinal cortex and hippocampus were dissected coronally, dehydrated and embedded in paraffin. The paraffin blocks were microtomed at 10 μm, mounted on Flex IHC adhesive microscope slides (Dako), and dried at 55 °C before storing. For neuropathological analysis, sections from all blocks were stained with anti-pTau (AT8), anti-Aβ (4G8) (Additional file 1, Table S1), and with the Gallyas and the Campbell-Switzer silver techniques for detection of neurofibrillary changes and amyloid deposits (43).

**Table 2.**
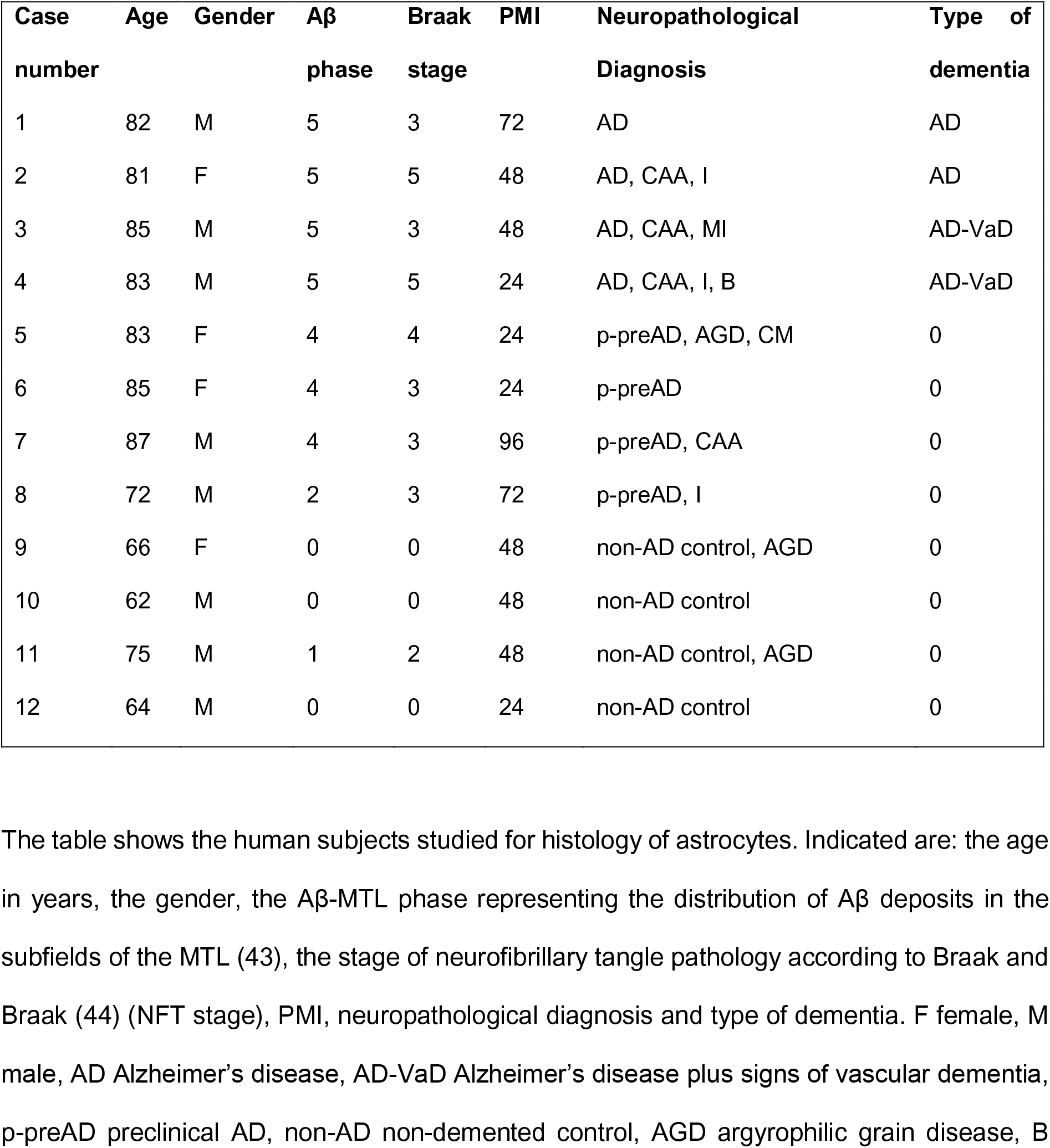
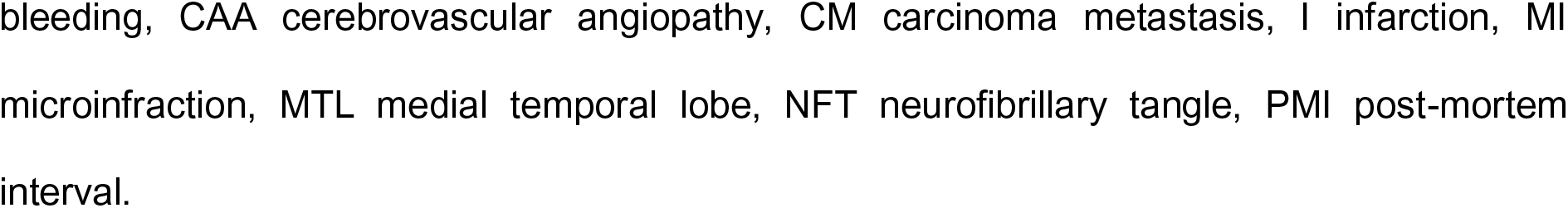
Details of the Human Cohort.

The post-mortem diagnosis of AD pathology was based upon the standardized clinico-pathological criteria, including the topographical distribution of Aβ plaques in the medial temporal lobe (AβMTL phase) based on Aβ immunohistochemistry (43), and the Braak neurofibrillary tangle (NFT) stage based on pTau immunohistochemistry (44). The study comprised 12 cases with an average age of 77 years and a female to male ratio of 4:8. The cases were divided in three groups based on the clinical and neuropathological diagnosis: (1) AD = high-intermediate degree of AD pathology and signs of cognitive decline during life (CDR ≥ 0.5); (2) p-preAD = cases with intermediate-low degrees of AD pathology lacking clinical signs of cognitive decline (CDR = 0); (3) non-AD = low-no pathological signs of AD pathology (CDR = 0).

### Immunohistochemistry and Immunofluorescence on Human Samples

The distribution of astrocytes and Aβ deposits was examined in human samples of the entorhinal cortex and hippocampus using immunohistochemical and immunofluorescence techniques. Immunohistochemical detection of Aβ deposits and astrocytes was performed after formic acid pretreatment. For double-labeling, a monoclonal anti-Aβ_17–24_ antibody (4G8, Additional file 1, Table S1) was subsequently combined with a polyclonal anti-GFAP (DAKO, Additional file 1, Table S1) as described previously (43). The anti-Aβ_17–24_ antibody was detected with biotinylated secondary antibodies and ABC, and visualized with 3,3’diaminobenzidine-HCl. After peroxidase blocking, the anti-GFAP was applied, detected with biotinylated secondary antibodies, and ABC, and visualized with the Vector peroxidase kit SG (blue staining). Microscopy analysis was performed using a light Leica DM2000 LED microscope (Leica Microsystems) and images were captured with a Leica DFC7000 T camera (Leica Microsystems).

For double-labeling immunofluorescence, sections were pre-treated as mentioned above and incubated with formic acid for 3 min, when required. Immunostainings were performed with an antibody cocktail and primary antibodies were detected with species-specific fluorescent-conjugated secondary antibodies (Additional file 1, Table S1). Images were captured via Nikon NIS-Elements software using a Nikon A1R laser scanning confocal system coupled to a Nikon Eclipse Ti inverted microscope (Nikon Instruments, Inc.). Acquired data were further processed using ImageJ software (National Institutes of Health).

## RESULTS

### Human iPSC-Derived Glial Progenitors Engraft the Mouse Brain and Differentiate into Astrocytes

To generate human-mouse astroglia chimeras, we differentiated human iPSCs (hiPSCs) into glial progenitor cells (hGPCs) *in vitro* (39) (Fig. 1a). After 44 days in culture, td-Tomato expressing hGPCs, which expressed several astroglia markers (Additional file 2, Figure S1b), were xenografted into the brains of newborn mice (Fig. 1a). We used transgenic Tg (Thy1-APPSw,Thy1-PSEN1*L166P) 21Jckr, also called APP/PS1-21 mice (40) crossed with immunodeficient NOD.CB17-Prkdc^scid^/J, further called NOD-SCID mice (41), to generate AD mice or wild-type (WT) littermates suitable for grafting experiments (31). We transplanted hiPSC lines from AD patients carrying the *APOE E4/E4* alleles and the corresponding corrected *APOE E3/E3* isogenic lines (Table 1).

Five months after transplantation, immunofluorescence (IF) analysis revealed engraftment of human cells throughout the forebrain (Fig. 1b, Additional file 2, Figure S1c). Human cells were identified based on the expression of the td-Tomato marker RFP and of the human nuclear antigen hNuclei. RFP+ cells infiltrate the cortex, corpus callosum and subcortical areas such as the hippocampus, striatum, thalamus or hypothalamus (Fig. 1c-e). Assessment of the engraftment capacity revealed considerable variation across cell lines (Additional file 2, Figure S1d): we show here examples of robust engraftment, with RFP+ cells both in clusters as well as integrated individually within the mouse brain (Fig. 1b, c), but these results were variable with often lower engraftment capacity at 5 months after transplantation (Additional file 2, Figure S1c, d). Variation was independent of the *APOE* genetic background or the patient (overview in Additional file 2, Figure S1d).

Further analyses revealed that at this stage, human RFP+ cells strongly express the astroglia markers GFAP, S100b, Vimentin and Aquaporin-4 (Fig. 1f-i), the latter largely concentrated at the astrocytic end-feet along the blood vessels (Figure 1i). Staining with human specific GFAP antibody (hGFAP), confirms the human origin of the cells (Additional file 3, Figure S2a). Quantification showed that 93% of the RFP+ hiPSC-cells express the astroglia marker GFAP (Fig. 1j) and 95% of the hNuclei+ hiPSC-cells co-express RFP (Fig. 1k). Thus, the RFP marker is not downregulated, and most of the transplanted cells indeed differentiated into human astroglia. This was further confirmed as no or only minimal expression (less than 3%) of neuronal or oligodendroglial markers was observed in RFP+ cells (Fig. 1l, Additional file 3, Figure S2b, c). No differences were observed between *APOE E4/E4* and *APOE E3/E3* lines (Additional file 3, Figure S2d-f). A subset of RFP+ cells identified by their distinct radial glia-like morphology and not expressing GFAP (Additional file 3, Figure S2g-i) often coexisted with RFP+ cells with more complex structures and expressing main astroglia markers. These cells are likely in a progenitor state which was also described previously (23,45).

### Transplanted iPSC-Derived Astrocytes Integrate Functionally Within the Mouse Brain

We assessed morphological and electrophysiological features of individual hiPSC-derived astrocytes in the chimeric brain. We observed hiPSC-astrocytes extending processes that terminated in end-feet contacting mouse host vasculature in the chimeric brains (Fig. 2a) similar to human astrocytes in the human brain (Fig. 2b). Moreover, hiPSC-astrocytes strongly expressed the gap-junction marker Connexin-43 in their processes (Fig. 2c). The gap junctions were functioning, as the Alexa488 dye loaded through the patch clamp pipette on RFP+ astrocytes diffused into neighboring mouse host cells (Fig. 2d-h). Electrophysiological analyses on acute brain slices of chimeric mice at 4-5 months showed that transplanted RFP+ astrocytes displayed properties resembling human astrocytes (46). Specifically, their non-excitable responses to stimulations with current injection in current clamp mode (Fig. 2i), resting membrane potentials (Fig. 2j), and linear current to voltage (I/V) curves (Fig. 2k). Human iPSC-astrocytes do not replace the endogenous murine astrocytes and both cell types are found in the chimeric mouse brains (Additional file 3, Figure S2j). These data reveal that the transplanted hiPSC-astrocytes are able to integrate functionally within the mouse host brain, show human-like physiological features and co-exist with endogenous mouse counterparts.

**Fig. 2.**
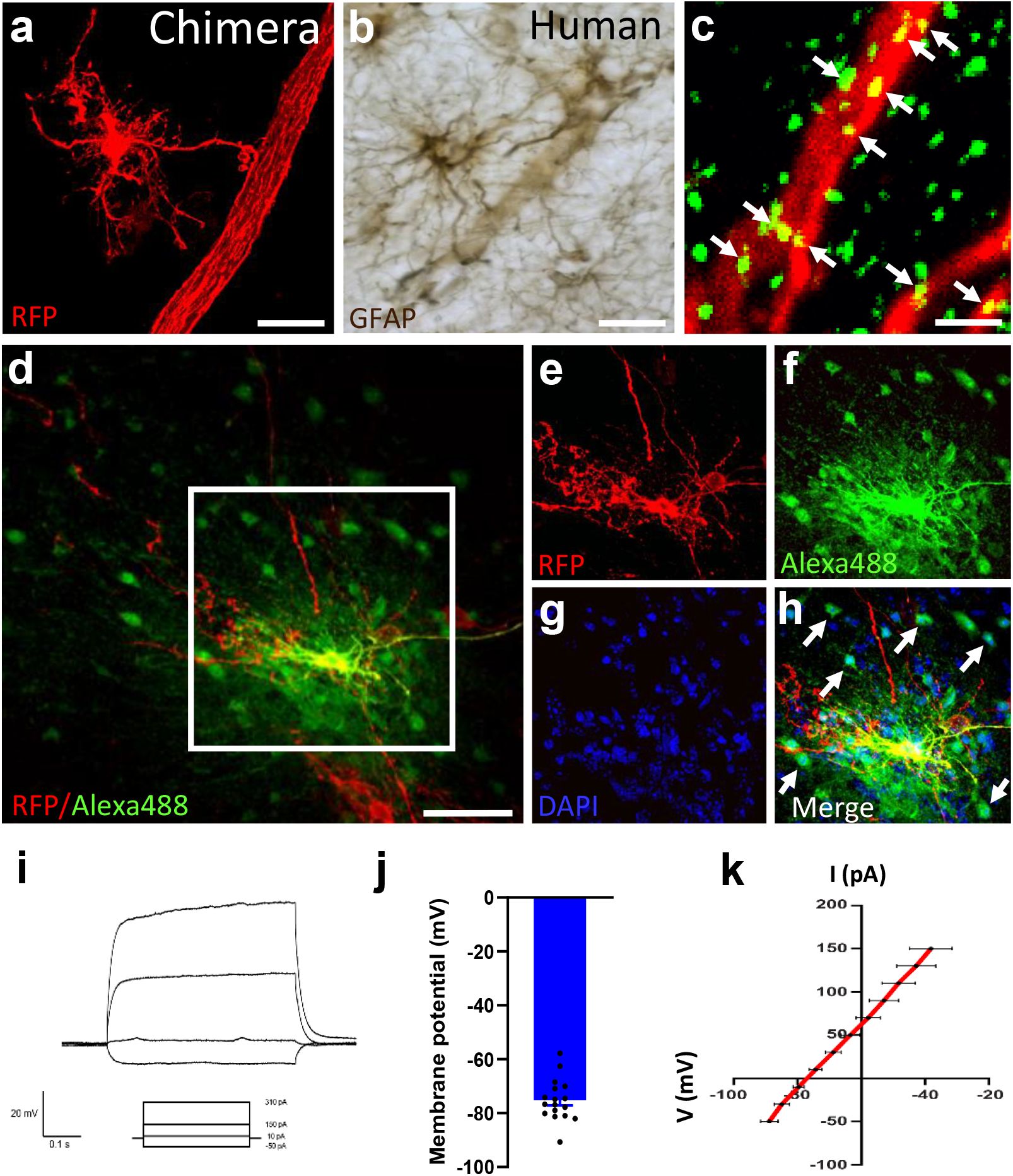
hiPSC-astrocytes integrate functionally within the mouse brain. **(a-b)** A xenografted hiPSC-astrocyte in the chimeric mouse brain (a, red) and a GFAP+ cortical astrocyte in the human brain (b, brown) contacting blood vessels with their end-feet. Scale bars: 25 μm. **(c)** hiPSC-astrocyte processes (RFP, red) express the gap junction marker Cx43 (green, arrows). Scale bar: 2 μm. **(d)** The gap-junction dye Alexa488 loaded on a hiPSC-astrocyte (RFP+, red) diffuses into RFP-neighboring host cells. Scale bar: 25 μm. **(e-h)** Enlarged views of the area selected in d. (e) RFP+ hiPSC-astrocyte, (f) Alexa488 dye, (g) Nuclei stained with DAPI, (h) Overlay. Arrows point to Alexa488+ RFP-host cells. **(i-k)** Representative traces of current injection steps of 20mV (i), resting membrane potentials (j) and current-voltage (I/V) curves (k) of hiPSC-astrocytes in the host brain (n=17 cells from 6 mice). Data are represented as mean ± SEM

### Human iPSC-Derived Astrocytes Acquire Human-Specific Morphologies and Features In Vivo

An advantage of low engraftment capacity is that it favors the assessment of morphological details of the transplanted astrocytes. Five months after transplantation, four main morphological subtypes of hiPSC-derived astrocytes were identified in the chimeric brains of the control animals. RFP+ interlaminar astrocytes were frequently observed in superficial layers of the cortex and close to the ventricles, with their small and round cell bodies near the pial surface and their long, unbranched and sometimes tortuous processes descending into deeper layers (Fig. 3a-c). Varicose-projection astrocytes were relatively sparse but easily identified by their bushy appearance and the presence of long processes with regularly spaced beads or varicosities (Fig. 3d, e). Protoplasmic astrocytes were found in deeper layers of the brain and showed the characteristic star-shaped morphology and shorter processes extending in all directions and often contacting the vasculature (Fig. 3f, g). Fibrous astrocytes were found in white matter tracts and presented the typical morphology with small soma and fine, straight and radially oriented processes (Fig. 3h-j). Interlaminar astrocytes were the most abundant subtype of hiPSC-astrocytes in the mouse brain, summing up to 62% of the RFP+ cells, and similar proportions of fibrous and protoplasmic astrocytes were found (16% and 13% of the RFP+ cells respectively). The varicose-projection astrocytes are the less frequent subtype, constituting 9% of RFP+ cells found in the host brain (Fig. 3k). Interestingly, we found the same astroglia subtypes in the human entorhinal cortex and white matter tracts of various control individuals (Table 2, subjects 10-12), when staining with the astrocyte marker GFAP: subpial interlaminar astrocytes with their soma in superficial layers of the cortex (molecular layer to pre-α) and long processes extending into deeper layers (Fig. 4a-c), protoplasmic (Fig. 4a, d) and varicose-projection astrocytes (Fig. 4e-f) in deeper layers of the cortex (pri-α to pri-γ), and fibrous astrocytes in white matter tracts (Fig. 4g-i). Of note, hiPSC-astrocytes covered about 15-fold larger areas than mouse astrocytes and displayed more complex structures (Fig. 3l-m, Additional file 3, Figure S2j). Thus, transplanted hiPSC-astrocytes were able to keep their intrinsic properties and develop in a cell-autonomous way adopting human-specific features and morphologies within the mouse host brain.

**Fig. 3.**
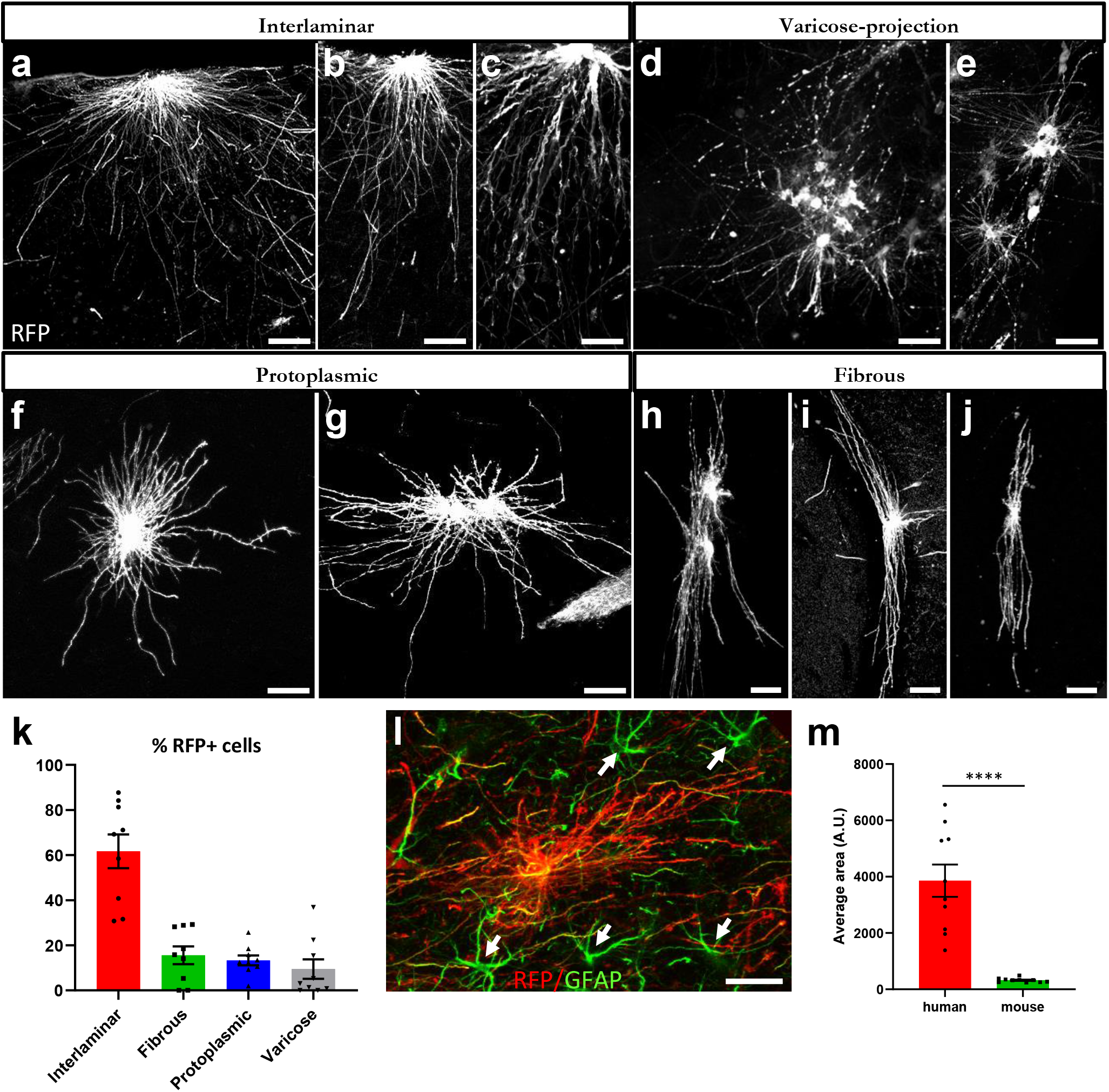
hiPSC-astrocytes recapitulate human morphological subtypes and retain human specific features within the mouse brain. **(a-j)** Representative images of RFP+ (white) interlaminar (a-c), varicose-projection (d-e), protoplasmic (f-g) and fibrous astrocytes (h-j) in the brain of wild-type mice five months after transplantation. Scale bars: 25 μm. **(k)** Histogram showing the percentage of RFP+ cells of each astroglial subtype on the mouse brain (n=9 mice). Data are represented as mean ± SEM. **(l)** Representative image showing mouse (green, arrows) and hiPSC-astrocytes (red) on a chimeric mouse brain five months after transplantation. Scale bar: 25 μm. **(m)** Histogram plotting the size of hiPSC-derived astrocytes vs mouse astrocytes on the host brain (n=12 mice). Data are represented as mean ± SEM, Student’s t test: ****p<0.0001

**Fig. 4.**
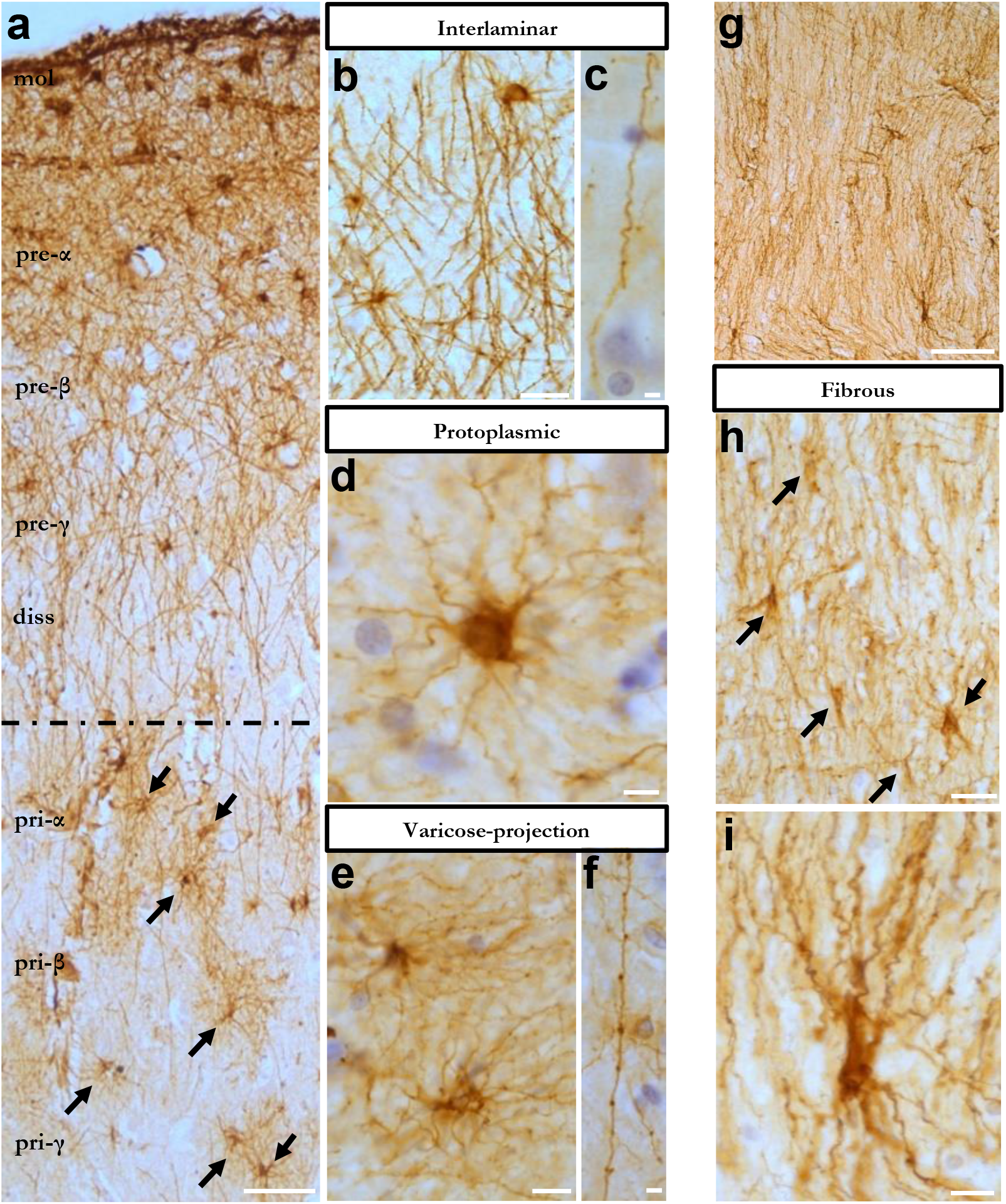
Four subtypes of morphologically defined GFAP+ astrocytes in the human entorhinal cortex and white matter. **(a)** Overview of human entorhinal cortex layers stained with GFAP (brown) to detect astrocytes. Layers molecular to lamina dissecans are mainly composed of subpial interlaminar astrocytes, while layers pri-α to pri-γ are rich in protoplasmic astrocytes (arrows). **(b-f)** Representative images of subpial interlaminar astrocytes (b) and their tortuous processes (c), protoplasmic astrocytes (d), varicose-projection astrocytes (e) and their beaded processes (f). **(g-i)** Overview of human white matter (g) and GFAP+ fibrous astrocytes (h-i). mol: molecular layer, diss: lamina dissecans. Scale bars: 50 μm in (a) and (g); 25 μm in (b) and (h); 10 μm in (c-f) and (i)

### Human Astroglia Display Differential Morphological Responses to Amyloid-β Plaques

Interestingly, transplanted hiPSC-astrocytes adopt three clearly distinct morphologies in the brains of chimeric AD mice five months after transplantation, when the Aβ load is high. Immunofluorescence analyses with RFP revealed that about 25% of the astrocytes became hypertrophic and showed thicker processes that surround Aβ deposits (Fig. 5a-c and 5a’-c’, Fig. 5g). 62% of the astrocytes seemed not to be morphologically affected at all, even when in close contact with the Aβ plaques (Fig. 5d, 5d’, 5g). Finally, about 13% of astrocytes showed atrophic features, displaying thinner processes that sometimes even looked degenerating (Fig. 5e-f and 5e’-f’, Fig. 5g). *APOE* E3/E3 or *APOE* E4/E4 genotype did not affect these proportions (Fig. 5h). Hypertrophic, atrophic and quiescent phenotypes were also found in human astrocytes in close proximity to Aβ deposits in the entorhinal cortex and hippocampus of patients with AD (Table 2, subjects 1-4), both by immunohistochemistry (Fig. 6) and immunofluorescence (Fig. 7).

**Fig. 5.**
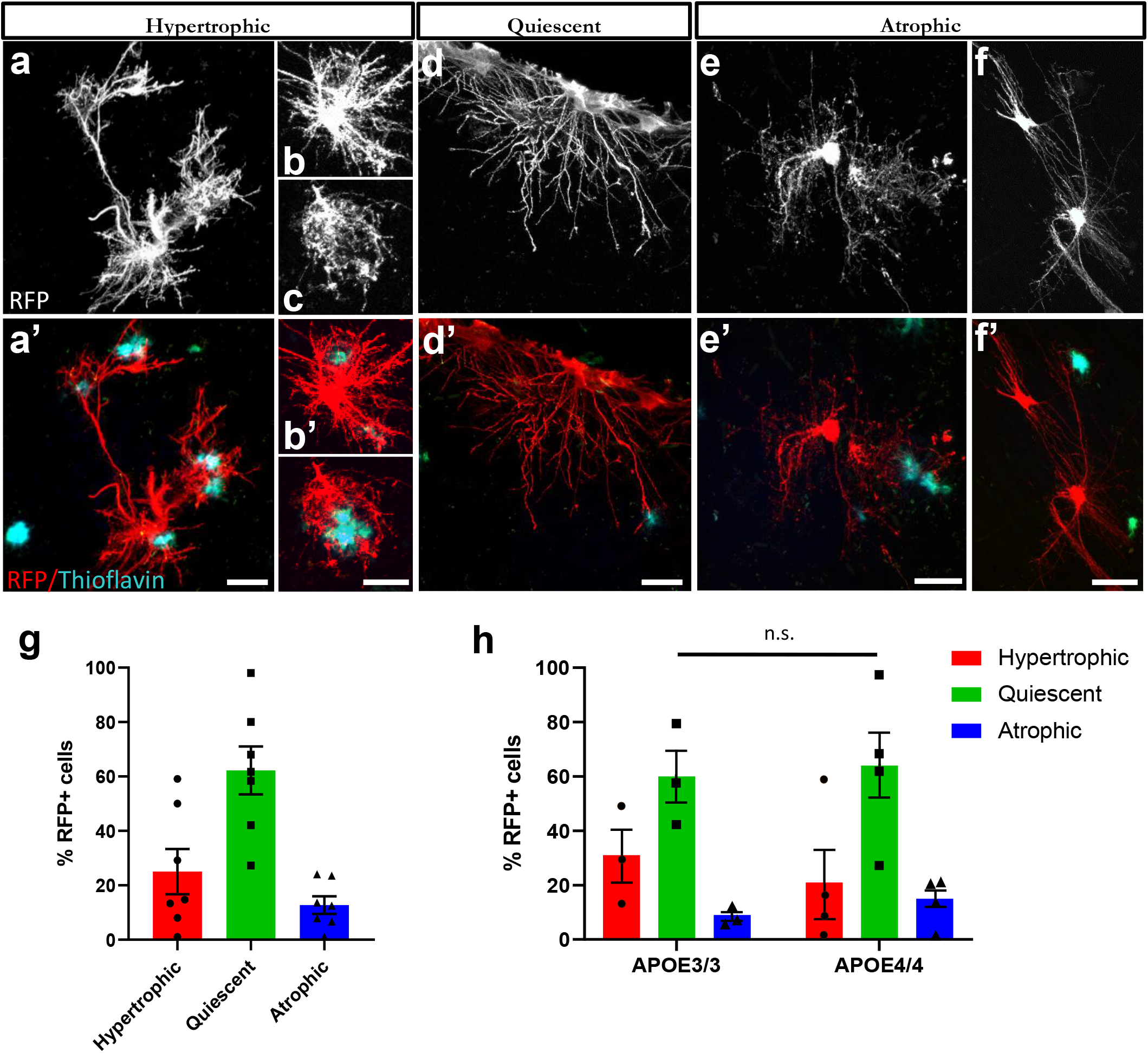
hiPSC-astrocytes show differential morphological responses to Aβ plaques within the chimeric mouse brain. **(a-f, a’-f’)** hiPSC-astrocytes (RFP+, red) exposed to Aβ plaques (Thioflavin, green) show hypertrophic (a-c, a’-c’), quiescent (d, d’) and atrophic (e-f, e’-f’) morphologies in AD chimeric mice five months after transplantation. Scale bars: 25 μm. **(g-h)** Percentage of hiPSC-astrocytes showing differential morphologies as a group (g, n=7 mice) and per ApoE genotype (h, n=3 mice for APOE3/3; n=4 mice for APOE4/4) five months post-transplantation. Data are represented as mean ± SEM, Chi-square test: n.s., non-significant

**Fig. 6.**
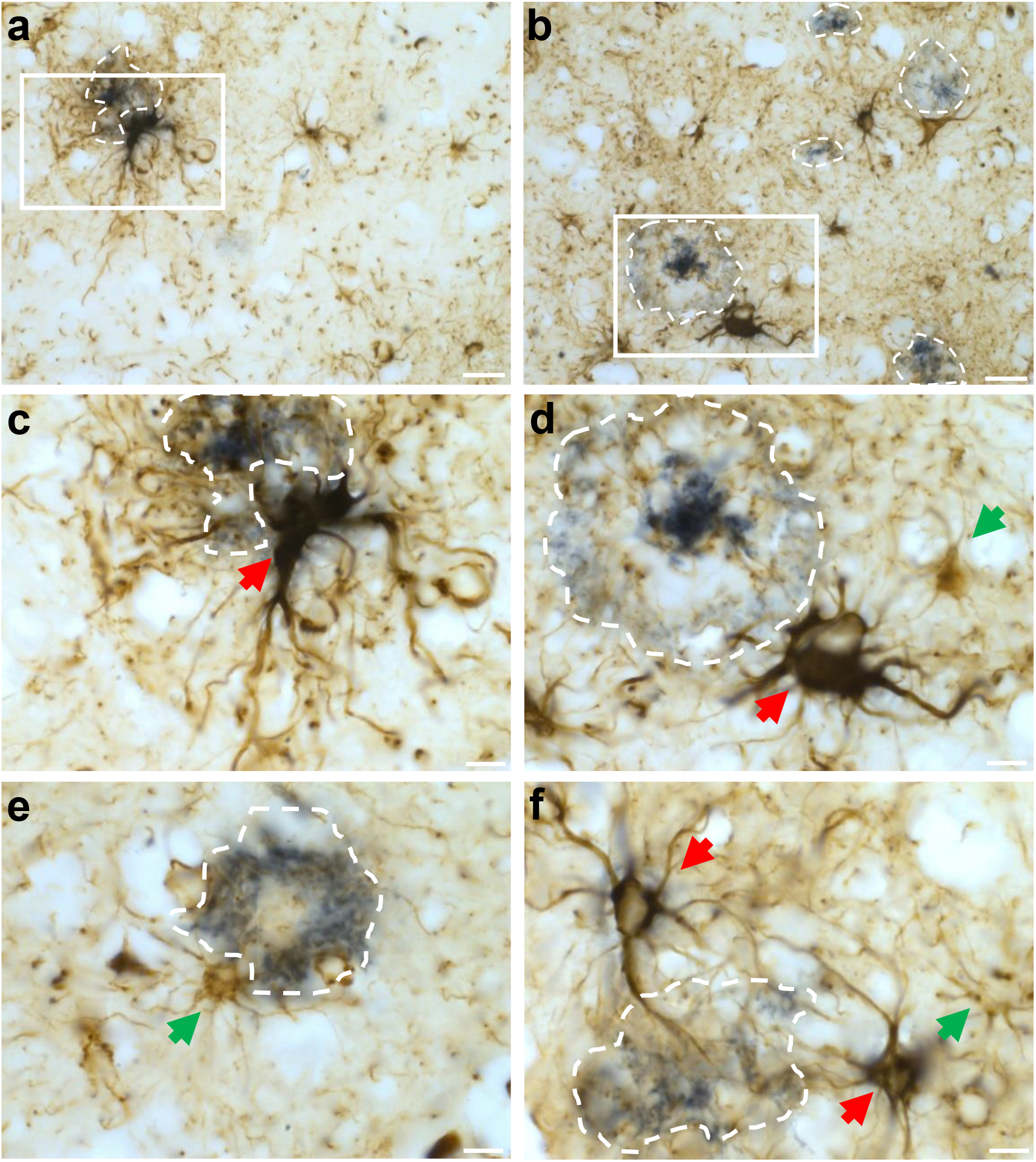
Astrocytes display differential responses to Aβ in the human AD-patient brain. **(a-f)** Representative immunohistochemistry images of GFAP+ astrocytes (brown) around amyloid-deposits (blue, dashed lines) in the cortex and hippocampus of AD-patient brains. **(a-d)** Overviews (a, b) and enlarged views (c, d) of the insets in a, b respectively. **(c-f)** GFAP+ hypertrophic (red arrows) and quiescent or atrophic (green arrows) astrocytes around amyloid-deposits. Scale bars: 25 μm in (a, b); 10 μm in (c-f)

**Fig. 7.**
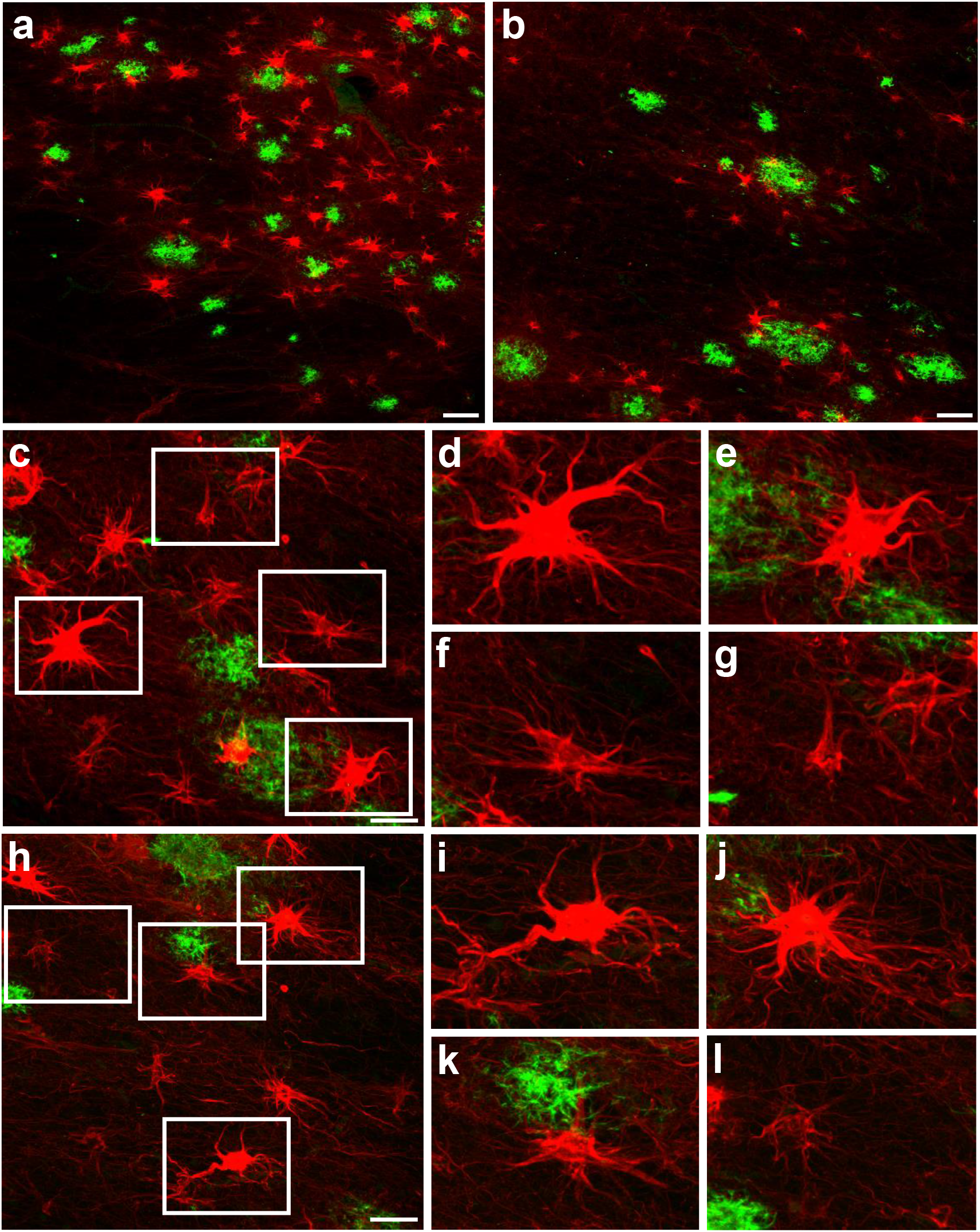
Hypertrophic, quiescent and atrophic astrocytes close to amyloid deposits in the human AD-patient brain. **(a-l)** Representative immunofluorescence images of GFAP+ astrocytes (red) around amyloid-deposits (4G8, green) in the cortex and hippocampus of AD patient brains. **(c-l)** GFAP+ astrocytes (red) show hypertrophic (d-e, i-j), quiescent (f, k) and atrophic (g, l) morphologies close to amyloid deposits. (d-g, i-l) Enlarged views of the insets in c and h respectively. Scale bars: 50 μm in (a, b) and 25 μm in (c, h)

In conclusion: engrafted hiPSC-astrocytes show differential morphological responses to Aβ plaques that resemble that of human astrocytes in AD patients’ brains. The potential of astrocytes to become hyper- or a-trophic, or remain in a quiescent state, does not seem to be influenced by the *APOE* genetic background.

## DISCUSSION

A major challenge to model astroglia function in AD is the difference between mouse and human astrocytes. A powerful approach to overcome this challenge is the use of hiPSC-derived astrocytes to generate chimeric mice.

We investigate in the current study the potential of such experiments using patient derived iPSC lines and isogenic counterparts (Table1). We include AD mice and control littermates.

We demonstrate integration of human glia into the mouse brain and differentiation of the majority of cells into four main subtypes of astrocytes expressing main astroglial markers and showing human-specific large, complex morphologies and electrophysiological properties. Additionally, hiPSC-astrocytes contact blood vessels and couple via gap-junctions with mouse cells, demonstrating functional integration in the host brain. In contrast to other glia chimeric models (35), we do not see replacement of the endogenous murine counterparts.

hiPSC-astrocytes respond robustly to Aβ pathology showing hypertrophic, atrophic or unaffected morphologies that are very similar to the morphological changes observed in astrocytes in AD patients’ brains (15–17). Such responses are not dependent on the *APOE* genetic background. Further work is however needed to understand whether the different *APOE* variants influence the molecular and functional states of human astrocytes surrounding Aβ plaques.

While human astrocytes were consistently detected in every injected brain, the number of engrafted cells varied largely from a few hundreds or thousands to >50,000 cells (Additional file 2, Figure S1d). This, combined with the difficulty of recovering the engrafted cells from the mouse brain for single-cell analysis, made further molecular analyses of the cellular phenotypes, unfortunately, not possible at this moment.

Others have also observed variations in transplantation efficiencies of hiPSC-derived microglia and neurons (47,48). While many successful reports on “glia” chimeric mice have been reported (23,34–36,49), these glia chimeras develop, in addition to human astrocytes, a large number of human NG2 cells and oligodendrocytes, whose relative ratios varied considerably across different brain regions and animals (23,34,49). This suggests that in these other experiments a different glia precursor state has been transplanted which maintains more ‘stem cell like’ properties allowing these cells to spread over the brain and to compete with mouse glia as shown before (49). We speculate that in our experimental conditions we have transplanted more differentiated cells which are closer to a final astrocyte phenotype and therefore not able to proliferate once they were injected in the brain of the host mice. It will now be critical to define the optimal window for transplantation of differentiating hiPSCs in order to maximize astrocyte colonization of the mouse brain. In other experiments we succeeded already to determine this for microglia using the Migrate protocol (Fattorelli et al, 2020). In the Migrate protocol there is a very critical window during the cell differentiation *in vitro* that results in 60-80% chimerism. One week longer in culture results in < 5% chimerism although the cells before transplantation look morphologically identical to the more efficiently transplanted ones. Other possible improvements would be the use of RAG2−/− mice which can be maintained for a much longer time period than the NOD-SCID mice we use here.

## CONCLUSIONS

In conclusion, despite some intrinsic limitations, the approach to transplant human astroglia into mouse brain to study astrocyte pathophysiology in AD is promising. We recapitulated here typical morphological responses of human astrocytes to amyloid plaques *in vivo*. Moreover, the combination of the model with isogenic *APOE* lines points out the potential use of this approach to analyze the impact of patient-derived and genetically modified astroglia on human CNS disease.

## LIST OF ABBREVIATIONS

5 M: 5 months of age
AD: Alzheimer’s disease
Aβ: amyloid-β
APOE: Apolipoprotein E
CDR: clinical dementia rating
DIV: days in vitro
EB: embryoid bodies
GPCs: glia progenitor cells
hiPSCs: human induced pluripotent stem cells
IF: immunofluorescence
IVC: individually ventilated cages
NFT: neurofibrillary tangles
NPCs: neural progenitor cells
PAM: protospacer adjacent motif
PMI: post-mortem interval
RFP: red fluorescent protein
RT: room temperature
SPF: specific pathogen free
ssODN: single-strand oligo-deoxynucleotide
Tzv: Thiazovivin
WT: Wild-type

## DECLARATIONS

### Ethics approval and consent to participate

All animal experiments were conducted according to protocols approved by the local Ethical Committee of Laboratory Animals of the KU Leuven (governmental licence LA1210591) following governmental and EU guidelines. All experiments conform to the relevant regulatory standards. The consent for reprogramming human somatic cells to hiPSCs was carried out on ESCRO protocol 19-04 at Mount Sinai (J.TCW.). The autopsies were performed with informed consent in accordance with the applicable laws in Belgium (UZ Leuven) and Germany (Ulm, Bonn and Offenbach). The use of human brain tissue samples for this study was approved by the ethical committees of Leuven University and UZ Leuven.

### Consent for publication

Not applicable.

### Availability of data and materials

The datasets used and/or analyzed during the current study are available from the corresponding authors on reasonable request.

### Authors’ contributions

AMA and BDS conceived the study and planned experiments. AMA, PP, JTCW, AS, SC, MAT, NC, and SM performed the experiments. All authors interpreted data. AMA and BDS wrote the first version of the manuscript. All authors contributed to and approved the final version.

## Acknowledgments

We thank Veronique Hendrickx and Jonas Verwaeren for help with the mouse colonies and Alicja Ronisz for technical assistance. Mouse experiments were supported by Inframouse (KU Leuven and VIB). Confocal microscopy was performed in the VIB Bio Imaging Core (LiMoNe and EMoNe facilities).

## REFERENCES

1. Ferrer I (2018) Astrogliopathy in Tauopathies. Neuroglia 1:126–150. doi: 10.3390/neuroglia1010010

2. Verkhratsky A, Nedergaard M (2018) Physiology of Astroglia. Physiol Rev 98:239–389. doi: 10.1152/physrev.00042.2016

3. Liddelow SA, Guttenplan KA, Clarke LE, Bennett FC, Bohlen CJ, Schirmer L, Bennett ML, Münch AE, Chung W-S, Peterson TC, Wilton DK, Frouin A, Napier BA, Panicker N, Kumar M, Buckwalter MS, Rowitch DH, Dawson VL, Dawson TM, Stevens B, Barres BA (2017) Neurotoxic reactive astrocytes are induced by activated microglia. Nature 541:481–487. doi: 10.1038/nature21029

4. Ouali Alami N, Schurr C, Olde Heuvel F, Tang L, Li Q, Tasdogan A, Kimbara A, Nettekoven M, Ottaviani G, Raposo C, Röver S, Rogers-Evans M, Rothenhäusler B, Ullmer C, Fingerle J, Grether U, Knuesel I, Boeckers TM, Ludolph A, Wirth T, Roselli F, Baumann B (2018) NF-κB activation in astrocytes drives a stage-specific beneficial neuroimmunological response in ALS. EMBO J. 37:e98697. doi: 10.15252/embj.201798697

5. Rothhammer V, Borucki DM, Tjon EC, Takenaka MC, Chao C, Ardura-fabregat A, Lima KA De, Gutiérrez-vázquez C, Hewson P, Staszewski O, Blain M, Healy L, Neziraj T, Borio M, Wheeler M, Dragin LL, Laplaud DA, Antel J, Alvarez JI, Prinz M, Quintana FJ (2018) Microglial control of astrocytes in response to microbial metabolites. Nature 557:724–728. doi: 10.1038/s41586-018-0119-x

6. Yun SP, Kam T, Panicker N, Kim S, Oh Y, Park J, Kwon S, Park YJ, Karuppagounder SS, Park H, Kim S, Oh N, Kim NA, Lee S, Brahmachari S, Mao X, Lee JH, Kumar M, An D, Kang S, Lee Y, Lee KC, Na DH, Kim D, Lee SH, Roschke VV, Liddelow SA, Mari Z, Barres BA, Dawson VL, Lee S (2018) Block of A1 astrocyte conversion by microglia is neuroprotective in models of Parkinson’s disease. Nat Med. 24:931–938. doi: 10.1038/s41591-018-0051-5

7. Wheeler MA, Clark IC, Tjon EC, Li Z, Zandee SEJ, Couturier CP, Watson BR, Scalisi G, Alkwai S, Rothhammer V, Rotem A, Heyman JA, Thaploo S, Sanmarco LM, Ragoussis J, Weitz DA, Petrecca K, Moffitt JR, Becher B, Antel JP, Prat A, Quintana FJ (2020) MAFG-driven astrocytes promote CNS inflammation. Nature 578:593–599. doi: 10.1038/s41586-020-1999-0

8. Arranz AM, De Strooper B (2019) The role of astroglia in Alzheimer’s disease: pathophysiology and clinical implications. Lancet Neurol 18:406–414. doi: 10.1016/S1474-4422(18)30490-3

9. Lambert JC, Ibrahim-Verbaas CA, Harold D, Naj AC, Sims R, Bellenguez C, et al. (2013) Meta-analysis of 74,046 individuals identifies 11 new susceptibility loci for Alzheimer’s disease. Nat Genet 45:1452–8. doi: 10.1038/ng.2802

10. Verheijen J, Sleegers K (2018) Understanding Alzheimer Disease at the Interface between Genetics and Transcriptomics. Trends Genet 34:434–447. doi: 10.1016/j.tig.2018.02.007

11. Zhang Y, Sloan SA, Clarke LE, Caneda C, Plaza CA, Blumenthal PD, Vogel H, Steinberg GK, Edwards MSB, Li G, Duncan JA, Cheshier SH, Shuer LM, Chang EF, Grant GA, Gephart MGH, Barres BA (2016) Purification and Characterization of Progenitor and Mature Human Astrocytes Reveals Transcriptional and Functional Differences with Mouse. Neuron 89:37–53. doi: 10.1016/j.neuron.2015.11.013

12. Thal DR, Schultz C, Dehghani F, Yamaguchi H, Braak H, Braak E (2000) Amyloid β-protein (Aβ)-containing astrocytes are located preferentially near N-terminal-truncated Aβ deposits in the human entorhinal cortex. Acta Neuropathol 100:608–617. doi: 10.1007/s004010000242

13. Thal DR (2012) The role of astrocytes in amyloid β-protein toxicity and clearance. Exp Neurol 236:1–5. doi: 10.1016/j.expneurol.2012.04.021

14. Mulder SD, Veerhuis R, Blankenstein MA, Nielsen HM (2012) The effect of amyloid associated proteins on the expression of genes involved in amyloid-β clearance by adult human astrocytes. Exp Neurol 233:373–379. doi: 10.1016/j.expneurol.2011.11.001

15. Pike CJ, Cummings BJ, Cotman CW (1995) Early association of reactive astrocytes with senile plaques in Alzheimer’s disease. Exp Neurol 132:172–179. doi: 10.1016/0014-4886(95)90022-5

16. Colombo JA, Quinn B, Puissant V (2002) Disruption of astroglial interlaminar processes in Alzheimer’s disease. Brain Res Bull 58:235–242. doi: 10.1016/S0361-9230(02)00785-2

17. Hsu ET, Gangolli M, Su S, Holleran L, Stein TD, Alvarez VE (2018) Astrocytic degeneration in chronic traumatic encephalopathy. Acta Neuropathol. 136:955–972. doi: 10.1007/s00401-018-1902-3

18. Orre M, Kamphuis W, Osborn LM, Jansen AHP, Kooijman L, Bossers K, Hol EM (2014) Isolation of glia from Alzheimer’s mice reveals inflammation and dysfunction. Neurobiol Aging. 35:2746–2760. doi: 10.1016/j.neurobiolaging.2014.06.004

19. Lian H, Yang L, Cole A, Sun L, Chiang ACA, Fowler SW, Shim DJ, Rodriguez-Rivera J, Taglialatela G, Jankowsky JL, Lu HC, Zheng H (2015) NFκB-Activated Astroglial Release of Complement C3 Compromises Neuronal Morphology and Function Associated with Alzheimer’s Disease. Neuron 85:101–115. doi: 10.1016/j.neuron.2014.11.018

20. Lian H, Litvinchuk A, Chiang AC-A, Aithmitti N, Jankowsky JL, Zheng H (2016) Astrocyte-Microglia Cross Talk through Complement Activation Modulates Amyloid Pathology in Mouse Models of Alzheimer’s Disease. J Neurosci 36:577–589. doi: 10.1523/JNEUROSCI.2117-15.2016

21. Diniz LP, Tortelli V, Matias XI, Morgado J, Be AP, Melo XHM, Seixas XGS, Alves-leon XS V, Souza XJM De, Ferreira XST, Felice XFG, De Gomes A (2017) Astrocyte Transforming Growth Factor Beta 1 Protects Synapses against Aβ Oligomers in Alzheimer’s Disease Model. Journal of Neuroscience 37:6797–6809. doi: 10.1523/JNEUROSCI.3351-16.2017

22. Oberheim NA, Takano T, Han X, He W, Lin JHC, Wang F, Xu Q, Wyatt JD, Pilcher W, Ojemann JG, Ransom BR, Goldman SA, Nedergaard M (2009) Uniquely Hominid Features of Adult Human Astrocytes. J Neurosci. 29:3276–87. doi: 10.1523/JNEUROSCI.4707-08.2009

23. Han X, Chen M, Wang F, Windrem M, Wang S, Shanz S, Xu Q, Oberheim NA, Bekar L, Betstadt S, Silva AJ, Takano T, Goldman SA, Nedergaard M (2013) Forebrain engraftment by human glial progenitor cells enhances synaptic plasticity and learning in adult mice. Cell Stem Cell 12(3):342–53. doi: 10.1016/j.stem.2012.12.015

24. Tarassishin L, Suh HS, Lee SC (2014) LPS and IL-1 differentially activate mouse and human astrocytes: Role of CD14. Glia 62:999–1013. doi: 10.1002/glia.22657

25. Lundin A, Delsing L, Clausen M, Ricchiuto P, Sanchez J, Sabirsh A, Ding M, Synnergren J, Zetterberg H, Brolén G, Hicks R, Herland A, Falk A (2018) Human iPS-Derived Astroglia from a Stable Neural Precursor State Show Improved Functionality Compared with Conventional Astrocytic Models. Stem Cell Reports 10:1030–1045. doi: 10.1016/j.stemcr.2018.01.021

26. Zhao J, Davis MD, Martens YA, Shinohara M, Graff-radford NR, Younkin SG, Wszolek ZK, Kanekiyo T, Bu G (2017) APOE e 4 / e 4 diminishes neurotrophic function of human iPSC-derived astrocytes. Hum. Mol. Genetics. 26:2690–2700. doi: 10.1093/hmg/ddx155

27. Oksanen M, Petersen AJ, Naumenko N, Puttonen K, Lehtonen Š, Gubert Olivé M, Shakirzyanova A, Leskelä S, Sarajärvi T, Viitanen M, Rinne JO, Hiltunen M, Haapasalo A, Giniatullin R, Tavi P, Zhang SC, Kanninen KM, Hämäläinen RH, Koistinaho J (2017) PSEN1 Mutant iPSC-Derived Model Reveals Severe Astrocyte Pathology in Alzheimer’s Disease. Stem Cell Reports 9:1885–1897. doi: 10.1016/j.stemcr.2017.10.016

28. Lin YT, Seo J, Gao F, Feldman HM, Wen HL, Penney J, Cam HP, Gjoneska E, Raja WK, Cheng J, Rueda R, Kritskiy O, Abdurrob F, Peng Z, Milo B, Yu CJ, Elmsaouri S, Dey D, Ko T, Yankner BA, Tsai LH (2018) APOE4 Causes Widespread Molecular and Cellular Alterations Associated with Alzheimer’s Disease Phenotypes in Human iPSC-Derived Brain Cell Types. Neuron 98:1141–1154.e7. doi: 10.1016/j.neuron.2018.05.008

29. Tcw J, Liang SA, Qian L, Pipalia NH, Chao MJ, Bertelsen SE, Kapoor M, Marcora E, Sikora E, Holtzman D, Maxfield FR, Zhang B, Wang M, Poon WW, Goate AM (2019) Cholesterol and Matrisome Pathways Dysregulated in Human APOE ɛ4 Glia. bioRxiv. 713362. doi: 10.2139/ssrn.3435267

30. Perriot S, Mathias A, Perriard G, Canales M, Jonkmans N, Merienne N, Meunier C, El Kassar L, Perrier AL, Laplaud DA, Schluep M, Déglon N, Du Pasquier R (2018) Human Induced Pluripotent Stem Cell-Derived Astrocytes Are Differentially Activated by Multiple Sclerosis-Associated Cytokines. Stem Cell Reports 11:1199–1210. doi: 10.1016/j.stemcr.2018.09.015

31. Espuny-Camacho I, Arranz AM, Fiers M, Snellinx A, Ando K, Munck S, Bonnefont J, Lambot L, Corthout N, Omodho L, Vanden Eynden E, Radaelli E, Tesseur I, Wray S, Ebneth A, Hardy J, Leroy K, Brion JP, Vanderhaeghen P, De Strooper B (2017) Hallmarks of Alzheimer’s Disease in Stem-Cell-Derived Human Neurons Transplanted into Mouse Brain. Neuron 93:1066–1081.e8. doi: 10.1016/j.neuron.2017.02.001

32. Mancuso R, Van Den Daele J, Fattorelli N, Wolfs L, Balusu S, Burton O, Liston A, Sierksma A, Fourne Y, Poovathingal S, Arranz-Mendiguren A, Sala Frigerio C, Claes C, Serneels L, Theys T, Perry VH, Verfaillie C, Fiers M, De Strooper B (2019) Stem-cell-derived human microglia transplanted in mouse brain to study human disease. Nat Neurosci. 22:2111–2116. doi: 10.1038/s41593-019-0525-x

33. Hasselmann J, Coburn MA, England W, Figueroa Velez DX, Kiani Shabestari S, Tu CH, McQuade A, Kolahdouzan M, Echeverria K, Claes C, Nakayama T, Azevedo R, Coufal NG, Han CZ, Cummings BJ, Davtyan H, Glass CK, Healy LM, Gandhi SP, Spitale RC, Blurton-Jones M (2019) Development of a Chimeric Model to Study and Manipulate Human Microglia In Vivo. Neuron 103:1016–1033.e10. doi: 10.1016/j.neuron.2019.07.002

34. Benraiss A, Wang S, Herrlinger S, Li X, Chandler-Militello D, Mauceri J, Burm HB, Toner M, Osipovitch M, Jim Xu Q, Ding F, Wang F, Kang N, Kang J, Curtin PC, Brunner D, Windrem MS, Munoz-Sanjuan I, Nedergaard M, Goldman SA (2016) Human glia can both induce and rescue aspects of disease phenotype in Huntington disease. Nat Commun. 7:11758. doi: 10.1038/ncomms11758

35. Windrem MS, Schanz SJ, Guo M, Tian GF, Washco V, Stanwood N, Rasband M, Roy NS, Nedergaard M, Havton LA, Wang S, Goldman SA (2008) Neonatal Chimerization with Human Glial Progenitor Cells Can Both Remyelinate and Rescue the Otherwise Lethally Hypomyelinated Shiverer Mouse. Cell Stem Cell. 2:553–565. doi: 10.1016/j.stem.2008.03.020

36. Windrem MS, Osipovitch M, Liu Z, Bates J, Chandler-Militello D, Zou L, Munir J, Schanz S, McCoy K, Miller RH, Wang S, Nedergaard M, Findling RL, Tesar PJ, Goldman SA (2017) Human iPSC Glial Mouse Chimeras Reveal Glial Contributions to Schizophrenia. Cell Stem Cell 21:195–208.e6. doi: 10.1016/j.stem.2017.06.012

37. Paquet D, Kwart D, Chen A, Sproul A, Jacob S, Teo S, Olsen KM, Gregg A, Noggle S, Tessier-Lavigne M (2016) Efficient introduction of specific homozygous and heterozygous mutations using CRISPR/Cas9. Nature 533:125–129. doi: 10.1038/nature17664

38. Bowles KR, Julia TCW, Qian L, Jadow BM, Goate AM (2019) Reduced variability of neural progenitor cells and improved purity of neuronal cultures using magnetic activated cell sorting. PLoS One 14:1–18. doi: 10.1371/journal.pone.0213374

39. Tcw J, Wang M, Pimenova AA, Bowles KR, Hartley BJ, Lacin E, Machlovi SI, Abdelaal R, Karch CM, Phatnani H, Slesinger PA, Zhang B, Goate AM, Brennand KJ (2017) An Efficient Platform for Astrocyte Differentiation from Human Induced Pluripotent Stem Cells. Stem Cell Reports 9:600–614. doi: 10.1016/j.stemcr.2017.06.018

40. Radde R, Bolmont T, Kaeser SA, Coomaraswamy J, Lindau D, Stoltze L, Calhoun ME, Jäggi F, Wolburg H, Gengler S, Haass C, Ghetti B, Czech C, Hölscher C, Mathews PM, Jucker M (2006) Abeta42-driven cerebral amyloidosis in transgenic mice reveals early and robust pathology. EMBO Rep 7:940–6. doi: 10.1038/sj.embor.7400784

41. Shultz LD, Schweitzer PA, Christianson SW, Gott B, Schweitzer IB, Tennent B, McKenna S, Mobraaten L, Rajan TV, Greiner DL (1995) Multiple defects in innate and adaptive immunologic function in NOD/LtSz-scid mice. J Immunol 154:180–91

42. Koper MJ, Van Schoor E, Ospitalieri S, Vandenberghe R, Vandenbulcke M, von Arnim CAF, Tousseyn T, Balusu S, De Strooper B, Thal DR (2020) Necrosome complex detected in granulovacuolar degeneration is associated with neuronal loss in Alzheimer’s disease. Acta Neuropathol 139:463–484. doi: 10.1007/s00401-019-02103-y

43. Thal DR, Rüb U, Schultz C, Sassin I, Ghebremedhin E, Del Tredici K, Braak E, Braak H (2000) Sequence of Aβ-protein deposition in the human medial temporal lobe. J Neuropathol Exp Neurol 59:733–748. doi: 10.1093/jnen/59.8.733

44. Braak H, Alafuzoff I, Arzberger T, Kretzschmar H, Tredici K (2006) Staging of Alzheimer disease-associated neurofibrillary pathology using paraffin sections and immunocytochemistry. Acta Neuropathol 112:389–404. doi: 10.1007/s00401-006-0127-z

45. Chen H, Qian K, Chen W, Hu B, Blackbourn LW, Du Z, Ma L, Liu H, Knobel KM, Ayala M, Zhang SC (2015) Human-derived neural progenitors functionally replace astrocytes in adult mice. J Clin Invest 125:1033–1042. doi: 10.1172/JCI69097

46. Sosunov AA, Wu X, Tsankova NM, Guilfoyle E, McKhann GM, Goldman JE (2014) Phenotypic heterogeneity and plasticity of isocortical and hippocampal astrocytes in the human brain. J Neurosci 34:2285–2298. doi: 10.1523/JNEUROSCI.4037-13.2014

47. Xu R, Li X, Boreland AJ, Posyton A, Kwan K, Hart RP, Jiang P (2020) Human iPSC-derived mature microglia retain their identity and functionally integrate in the chimeric mouse brain. Nat Commun 11: 1577. doi: 10.1038/s41467-020-15411-9

48. Kirkeby A, Nolbrant S, Tiklova K, Heuer A, Kee N, Cardoso T, Ottosson DR, Lelos MJ, Rifes P, Dunnett SB, Grealish S, Perlmann T, Parmar M (2017) Predictive Markers Guide Differentiation to Improve Graft Outcome in Clinical Translation of hESC-Based Therapy for Parkinson’s Disease. Cell Stem Cell 20:135–148. doi: 10.1016/j.stem.2016.09.004

49. Windrem MS, Schanz SJ, Morrow C, Munir J, Chandler-Militello D, Wang S, Goldman SA (2014) A Competitive Advantage by Neonatally Engrafted Human Glial Progenitors Yields Mice Whose Brains Are Chimeric for Human Glia. J Neurosci. 34:16153–16161. doi: 10.1523/JNEUROSCI.1510-14.2014

