## Additional file 1. Table S1 for "Human iPSC-derived astrocytes transplanted into the mouse brain display three morphological responses to amyloid-β plaques"

**Table S1. Antibodies used in this study.** The table summarizes information about supplier company, catalog number and concentration of use

| **Antibody** | **Company** | **Cat #** | **Concentration** |
| --- | --- | --- | --- |
| 4G8 | BioLegend | 800703 | 1/5000 |
| APC | Millipore | OP80 | 1/200 |
| AQP4  AT8 | Alomone Labs  Thermo Fisher Scientific | AQP-004  MN1020 | 1/300  1/100 |
| Cx43 | Santa Cruz | sc-271837 | 1/200 |
| EAAT1 (GLAST; ACSA-1)  FoxP2 | Miltenyi Biotec  Abcam | 130-095-814  Ab16046 | 1/200  1/100 |
| GFAP | Synaptic Systems | 173 004 | 1/1000 |
| GFAP | DAKO | Z033401-2 | 1/1000 |
| Glutamine Synthetase | Millipore | MAB302 | 1/100 |
| human GFAP | BioLegend | 837202 | 1/200 |
| human Nuclear Antigen  Nestin | Millipore  Abcam | MAB1281  Ab22035 | 1/100  1/200 |
| NeuN  PAX6 | Synaptic Systems  Abcam | 266 004  Ab5790 | 1/300  1/100 |
| RFP | Rockland | 600-401-379 | 1/1000 |
| RFP | Rockland | 200-301-379 | 1/1000 |
| S100b  SOX2 | Abcam  Cell Signaling | ab52642  3579S | 1/500  1/300 |
| STEM123 | Takara | Y40420 | 1/100 |
| Vimentin | BD Pharmingen | 550513 | 1/100 |
| Anti-Mouse 488 | Invitrogen | A21202 | 1/500 |
| Anti-Mouse 594 | Invitrogen | A21203 | 1/500 |
| Anti-Mouse 647 | Invitrogen | A31571 | 1/500 |
| Anti-Mouse biotinylated | Vector Laboratories | BA-9200 | 1/250 |
| Anti Mouse HRP | DAKO | P 0447 | 1/200 |
| Anti-Rabbit 488 | Invitrogen | A21206 | 1/500 |
| Anti-Rabbit 594 | Invitrogen | A21207 | 1/500 |
| Anti-Rabbit biotinlylated | Vector Laboratories | BA-1000 | 1/250 |
| Anti-Guinea Pig 647 | Jackson ImmunoResearch | 706-175-148 | 1/500 |
