## Additional file 2. Figure S2 for "Human iPSC-derived astrocytes transplanted into the mouse brain display three morphological responses to amyloid-β plaques"

**Figure S1 Characterization of hiPSC lines, derived glial progenitors and engraftment capacity. (a)** Karyotype analysis of the hiPSC lines used for transplantation showing no genomic alterations. **(b)** hiPSC-derived glial progenitors at 44 days in vitro express main astroglial markers. Scale bar: 50 µm. **(c)** Coronal sections stained with RFP show the distribution of xenografted hiPSC-derived astrocytes on a chimeric mouse brain at five months post-transplantation. Scale bar: 200 µm. **(d)** Assessment of engraftment capacity of trasplanted hiPSC-derived astrocytes: * indicates less than 5,000 RFP+ cells, ** between 5,000 and 25,000 RFP+ cells, and *** more than 25,000 RFP+ cells in the chimeric mouse brain

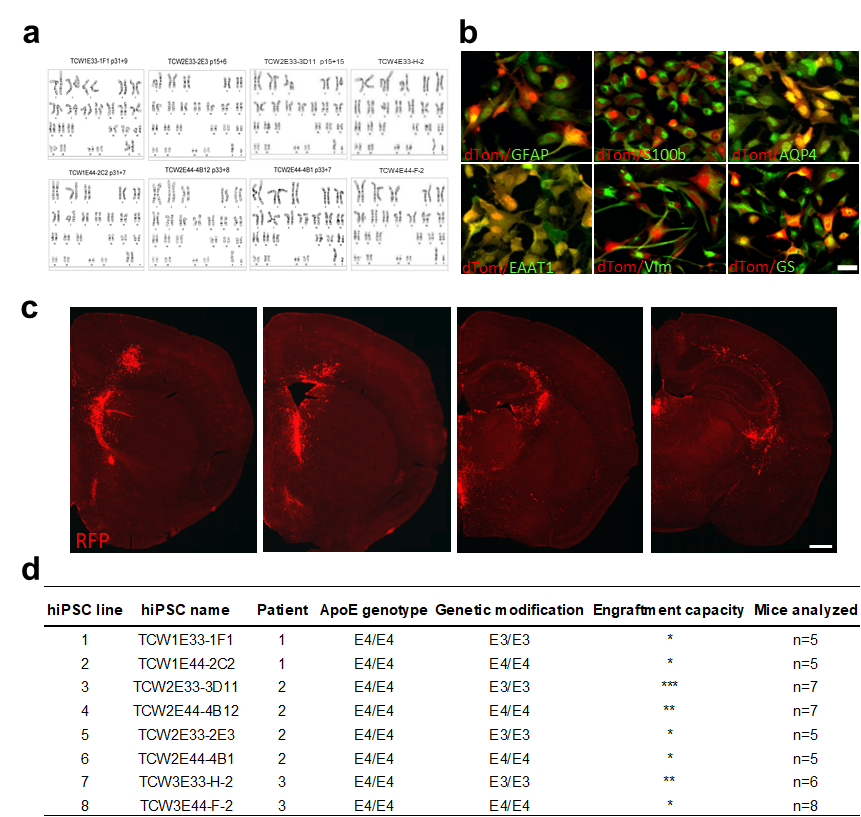
