## Additional file 3. Figure S3 for "Human iPSC-derived astrocytes transplanted into the mouse brain display three morphological responses to amyloid-β plaques"

**Figure S2 Characterization of hiPSC derived glia in vivo. (a-c)** Five months after transplantation, hiPSC-glia (RFP+, red) express human GFAP (a; hGFAP, green) but not the neuronal marker NeuN (b; green) or the oligodendrocyte marker APC (c; green). Scale bars: 25 µm. **(d-f)** Percentage of RFP+ cells expressing GFAP (d; n=7 mice for ApoE33; n=7 mice for ApoE44), NeuN (e; n=6 mice for ApoE33; n=3 mice for ApoE44) or APC (f; n=6 mice for ApoE33; n=3 mice for ApoE44). Data are represented as mean ± SEM, Student’s t test: n.s., non-significant. **(g-i)** Overview (g) and representative images (h-i) of RFP+ and GFAP- progenitor cells showing morphological features of radial glia. Scale bars: 25 µm. **(j)** hiPSC-astrocytes (RFP+ GFAP+) and mouse astrocytes (RFP- GFAP+) coexist within chimeric brains. Scale bar: 25 µm

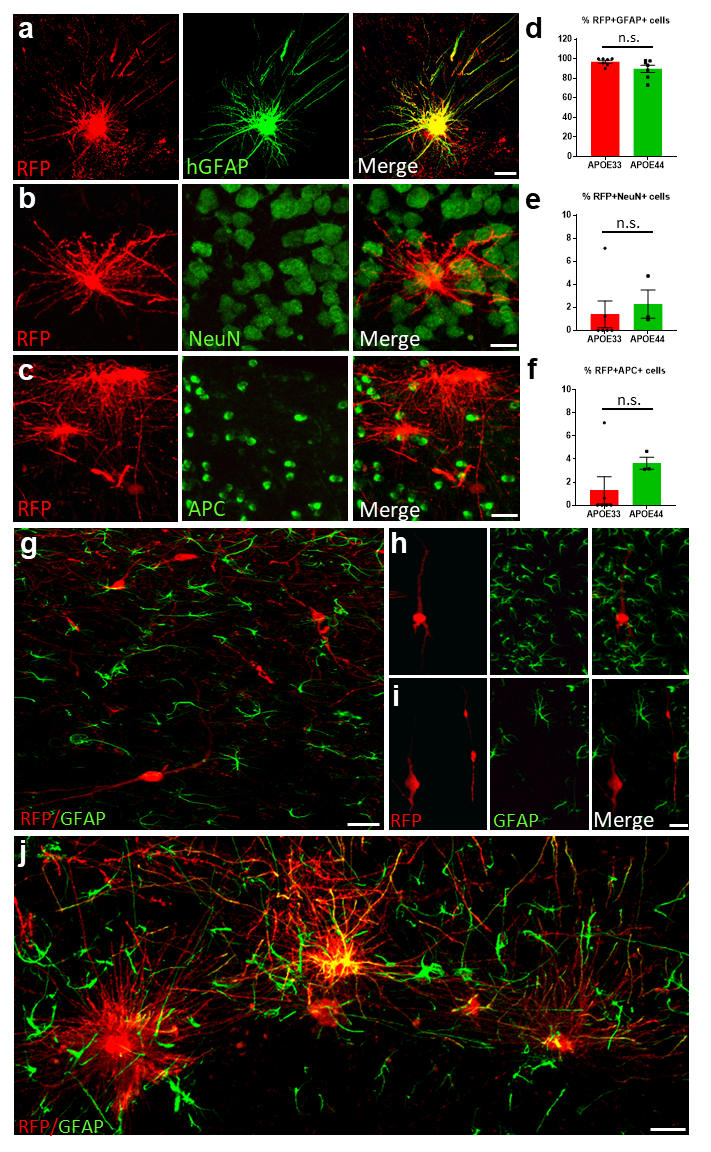
